# Deciphering the role of the nanoscale clustering of activating and costimulatory antibodies on T cell activation

**DOI:** 10.1101/2024.09.28.615509

**Authors:** Guillaume Le Saux, Piotr Nowakowski, Esti Toledo, Sivan Tzdaka, Abed Al-Kader Yassin, Shagufta Naaz, Yuval Segal, Brit Maman, Angel Porgador, Ana-Sunčana Smith, Mark Schvartzman

## Abstract

In a joint experimental and a modeling effort, we explored the regulation of T cell activation and cytotoxicity by the nanoscale clustering and surface density of activating and costimulatory antibodies. Specifically, we simulated T cells on nanolithographically patterned arrays of clusters of these antibodies, systematically varying the cluster size from intermediate to large and overall antibody density. We found that T-cell activation correlated more with global antibody density than with cluster size, such that at low density arrays were inefficient in activation, while high density arrays saturated the signal. However, when T-cells were exposed to patterns of low global densities but small, very dense clusters, full activation of T cells was achieved. These results could be rationalized using the membrane-fluctuation-model that integrates the cooperative effects between bonds with mechanical feedback from the cell activation. This insight into the spatial organization of activating ligands provides an important understanding of the mechanism of T cell activation and allows for the design of more effective T cell activation platforms for immunotherapy.

---

Adoptive immunotherapy, based on *ex vivo* engineered T cells, is a revolutionary approach to cancer treatment^1^. The engineered T cells are designed to recognize specific tumor-associated antigens, allowing them to effectively identify and eliminate cancer cells. Ongoing research, encompassing more than 1000 clinical trials, continues to refine and broaden the application of this transformative therapeutic approach. This therapeutic strategy involves isolation and subsequent *ex vivo* modification and expansion of a patient’s own T cells outside before their infusion to target cancer cells. The process begins with the extraction of T cells from the patient’s blood. This is followed by activation of cells and genetic engineering techniques to enhance their cancer-fighting capabilities, often through the introduction of chimeric antigen receptors (CARs)^2^. These modified T cells are then expanded in large quantities in the laboratory before being infused back into the patient.

The key step in the T cell engineering is their activation, which is critical for proliferation, differentiation, and effective transduction with synthetic receptors specific for cancer markers. The *in vivo* activation process by which T cells are primed by antigen-presenting-or target-cells^3,4^ requires three essential stimuli: (*i*) triggering T Cell Receptor (TCR) signaling with peptide-loaded major histocompatibility complexes (MHC), (*ii*) triggering CD28 co-stimulatory signaling with its cognate ligands, *e*.*g*., CD80, and (*iii*) providing cytokine to T cells.

The *ex vivo* activation process is largely inspired by the mechanism by which T cells activate *in vivo*, but now, natural ligands are commonly replaced with antibodies against costimulatory CD28, and CD3 protein that form a complex with TCR. While these antibodies can be supplied to T cells in a soluble manner, effective activation requires them to be physically tethered to a supporting surface, similar to natural ligands being tethered to the target cell membrane. According to the existing models of T cell activation, this tethering is essential for the spatial exclusion of large phosphatase molecules from the binding region to shift the kinase–phosphatase balance towards kinase-triggered activation signaling^5,6^, as well as for conformational changes in the bound receptors^7^. These local contacts are also believed to facilitate more binding of ligand–receptor pairs around the already bound ligand–receptor, leading to the formation of receptor clusters sized from a few tens to a few hundreds of nanometers^8–12^. Still, the role of this clustering is debated, and in particular, it is unclear whether this clustering is a precondition or a consequence of the receptor triggering and activating signaling.

Although both *in vivo* and *ex vivo* triggering of receptors in T cells target the same signaling pathways, they differ strikingly in several aspects. *In vivo*, the relatively low ligand–receptor affinity and high on–off ligand–receptor binding rate allow for a serial triggering mechanism by which one single ligand can trigger multiple TCRs and even activate entire T cell^13^. Such serial triggering is not possible with antibodies, whose binding affinity is an order of magnitude higher than that of natural ligands. Thus, T cell activation requires a relatively large number of antibodies, as compared to number of ligands in pMHC-based activation^14^. Furthermore, *in vivo*, both the ligands and receptors are mobile within the cell membrane, allowing a highly dynamic formation and arrangement of clusters even at very low ligand concentrations^15^, creating optimal conditions for activation^16^. On the contrary, in typical *ex vivo* activation platforms, the antibodies are statically connected to the surface, fixing the position of the bonded receptors. Examples of such platforms are antibody-coated microbeads that are the current golden standard in immunotherapeutic T cell production, as well as various recently proposed alternatives to these beads^17–20^. Remarkably, the spatial distribution of the antibodies that tether their cognate receptors of T cells was not considered as a parameter in the design of T cell activating materials.

Besides the simplicity of production, the reason for utilizing only systems with homogeneous distribution of ligands or antibodies is the lack of consensus about the role of clustering in T cell activation (for a recent review see Ref. ^21^). So far, studies using self-assembled hexagonal arrays of continuously arranged antibodies clearly showed that the density of ligands or antibodies is crucial for T cell activation^22,23^. However, these systems could not elucidate the difference between the role of overall density and organization. More sophisticated antibody arrays, with independently varied clustering and density, were shown to spatially control receptor binding, but no impact of TCR clustering on spreading properties of T cells could be determined.^23^ However, effects of antibody density and clustering on T cell activation and immune response were not evaluated. Furthermore, all these studies were focused solely on anti-CD3 antibodies, completely ignoring the essential costimulatory CD28 signaling.

In this work, we circumvent these issues in a study of the activation of T cells, which focuses on the effect of the overall surface density and organization of the activating and inhibitory antibodies. The antibodies are confined into clusters whose size ranges from 100 nm to 1 μm. The clusters themselves are arranged into periodic arrays with a periodicity that is double the cluster diameter. These cluster sizes and periodicity mimic the ligand–receptor nano-clusters observed at the physiological interface between T cells and target cells^24^. Besides local density (*i*.*e*., density of antibodies within each cluster), we also vary the global density of antibodies (*i*.*e*., overall number of antibodies per unit area of the system) by diluting them in mock antibodies with no specific binding. We stimulate primary T cells on these arrays and find that their activation and cytotoxic function directly correlate with the global antibody density but not with the cluster size. We then engineer a new array in which the global antibody density is kept at its lowest value (low activation if homogeneously distributed), but the receptors are grouped in small dense clusters. In these extreme, but still physiological, conditions we find a strong boost in T cell activation. We can rationalize these findings using the recently proposed Membrane Fluctuation Model (MFM) that couples the spatial distribution of ligands to the cell activity^25^. We are thus able to reveal the separate effects of the (*i*) antibody confinement within the clusters and (*ii*) the global surface density. We can now show that the nanoscale clustering of the activating and costimulatory antibodies is an important regulator of T cell response—an insight that is critical for our understanding of immunity and designing new biomaterials for immunomodulation.

## Assay design and fabrication

Nanoclusters of anti-CD3/anti-CD28 were fabricated using nanosphere lithography, which is an ideal approach to produce large area nanoscale patterns that mimic extracellular environment^26^. First, polystyrene nanospheres were assembled on a silicon surface into a closely packed polycrystalline monolayer. For 200 nm nanospheres, the assembly was done using the Langmuir–Blodgett method. For larger particles, the so-called “dry method” was employed, by which the particles were first rubbed between two Polydimethylsiloxane (PDMS) surfaces to form a closely packed monolayer, which was then mechanically transferred to silicon coated with a thin film of Polyethyleneimine (PEI)^27^. In all cases, the assembled particles were etched by reactive ion etching to half of their original diameter. A thin layer of gold was then evaporated through the formed nanosphere mask. In the next step, nanosphere lift-off was used to obtain circular discs of silicon surface within a continuous gold mesh (Fig. S1). These silicon discs were used as confinements for the antibodies, due to multiple straight-forward strategies to selectively functionalize Silicon with cell-interacting biomolecules^28^. To that end, the discs were functionalized with a homogeneous mix of anti-CD3 and anti-CD28. For that purpose, the disks were first coated with a monolayer of (3-aminopropyl)triethoxysilane (APTES), followed by the attachment of polyethylene glycol (PEG)-carboxylic acid terminated with biotin, to which biotinylated anti-CD3 and anti-CD28 were attached through a neutravidin bridge. The gold surrounding of the silicon discs was passivated with thiol-polyethylene glycol to prevent non-specific binding. The site-specificity of the biofunctionalization was examined using fluorescently tagged neutravidin. The quality of the layer was assessed using confocal fluorescence microscopy where fluorescent patterns with a geometry that corresponds to the fabricated array of silicon discs, could be observed (inset of Fig. 1b).

**Figure 1:**
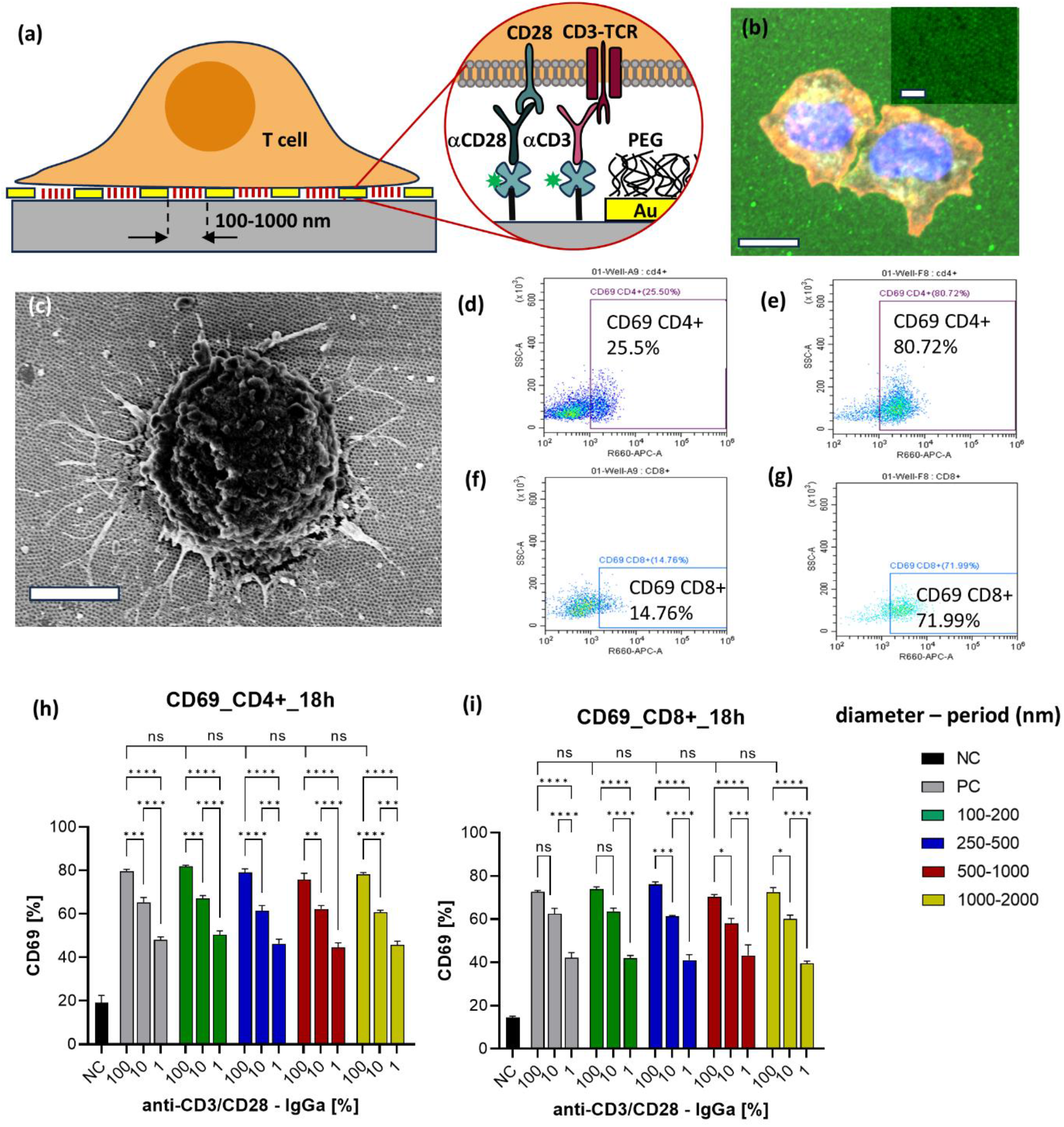
Activation of T-cell measured through CD69 expression levels. (a) Schematic plot of nano-engineered platform for the control of the receptor clustering with the patterned antibodies. (b) Fluorescent microscopy of T cells spread on an array of antibody clusters. Avidin used for binding the antibodies to Silicon surfaces was tagged with green, and the cells were tagged with red phalloidin for cytoskeleton and blue DAPI for nucleus. Scale bar is 5 μm. Inset: High resolution image of clusters visualized through the tagged neutravidin. Scale bar is 1 μm. (c) Scanning electron micrograph of T cell spread on the nanoarray, Scale bar : 2 microns (d) and (e) FC analysis of T CD69 expression by CD4+ T cells stimulated on negative control and 100 nm clusters with no dilution, respectively. (f) and (g) Flow Cytometry (FC) analysis of T CD69 expression by CD8+ T cells stimulated on negative control and 100 nm clusters with no dilution, respectively. (h) and (i) CD69 expression for CD4+ T cells and CD8+ T cells stimulated on various antibody clusters and dilutions. NC and PC stand for negative and positive controls, respectively. The statistical analysis was performed with Tukey’s multiple-comparison tests using the GraphPad Prism software. * *p <* 0.05, ** *p <* 0.01, *** *p <* 0.001, **** *p <* 0.0001, ns: not significant.

To examine the effect of the antibody surface density independently of the cluster size, “diluted” arrays were also produced, in which α-CD3 and α-CD28 were mixed with human immunoglobulin G2 (IGG2). Specifically, we used ratios of 1:10 (α-CD3/α-CD28:IGG2) and 1:100 (α-CD3/α-CD28:IGG2), termed here as 10% dilution and 1% dilution, respectively. These arrays are denoted hereafter by *cluster diameter (nm)/periodicity (nm)/ dilution (%)*. Finally, surfaces continuously covered with antibodies—undiluted and diluted with IGG2, and surfaces lacking antibodies, were used as positive and negative controls, respectively.

Peripheral Blood Mononuclear Cells (PBMCs) used as a source of T cells were isolated from the blood of a healthy donor. Cells were seeded onto the patterned surfaces, and mounted to the bottom of wells in a 96-well plate. Following the stimulations, the cells were stained for CD3, CD4, CD8, and with the activation marker CD69, whose expression was assessed by flow cytometry. Also, the cells were fixed and stained for the cytoskeleton with phalloidin and for the nucleus with DAPI for microscopic visualization. This visualization showed that cells spread onto the activating surfaces and produced tight contacts with multiple clusters (Fig. 1b). The morphology of the spread cells was examined in more detail using scanning electron microscopy (Fig. 1c). Here, clear nanometric protrusions were observed. These protrusions clearly resembled, in their size and shape, the physiological “microvilli” with which T cells probe their surroundings *in vivo*^29,30^.

## Effect of aCD3/aC28 distribution on T cell activation

The activation was assessed separately for CD4+ and CD8+ T cells through the expression of CD69 (Fig. 1d–i), after 18 hours of activation. The percentages of CD69 positive cells are shown in Fig. 1f– i for different clustering configurations, and global densities of activating and costimulatory antibodies. The first and foremost conclusion from this data is that the activation degree is not affected by the size of the antibody cluster within the probed cluster sizes and arrangements. The activation is, however, strikingly dependent on the overall density of the activating and costimulatory antibodies. This dependence has repeating patterns for both the T cell subsets and activation times, consistently with results on activating T cells using exclusively anti-CD3^21^.

Both CD4+ and CD8+ T cells reach similarly high activation levels after 18 hours, resulting in around 80% of CD69-positive cells for densely packed antibodies within the cluster. Interestingly, the same level of activation was produced by positive control surfaces with continuously arranged antibodies. Surface dilution of antibodies in IGG2, however, gradually reduced the activation level, reaching around 40% activation for 1% dilution. This was still higher than the activation produced by the negative control surface. In the latter case, the activation could be attributed to the tonic signaling produced by the surface lacking any activating/costimulatory antibodies. Similar trends were obtained for positive control surfaces with the continuous coating by the antibodies and similar degrees of dilutions, further emphasizing the role of overall density on these patterns.

In the controls, the continuous antibody surfaces not only produced activation similar to the surfaces patterned with clusters but also showed the same dependence of the activation on the global antibody density. Finally, it should be noted that while the CD69 signal is analyzed in terms of the Mean Fluorescence Intensity per cell (MFI) per sample, its dependence on the clustering and global density of antibody precisely mirrors that obtained for the percentage of CD69 positive cells (Fig. S2). Overall, these experimental findings indicate that the global antibody density rather than their clustering is the main regulator of T cell activation associated with CD69, for the cluster geometries probed here.

## Effect of αCD3/αC28 distribution on T cell cytotoxic activity

We, furthermore, assessed the cytotoxic activity of T cells using CD107a—a marker expressed on the surface of cytotoxic T cells during the degranulation of lytic granules. Here, the activation lasted for four hours, according to the standard protocol of the CD107a assay. Accordingly, the tagged anti-CD107a was added to the solution at the beginning of the activation, to allow for the detection of the accumulation of CD107α on the cell membrane throughout the entire activation process. CD4+ T cells showed negligible degranulation, which is attributed to their non-cytotoxic nature. To the contrary, we obtained a relatively high degree of CD8+CD107α+ T cells already after four hours, as anticipated (Fig. 2). The fraction of CD8+CD107α+ T cells (Fig. 2f) and the expression CD107α+ determined from the MFI (Fig. S3) mirrors the trend observed for CD69: there is no statistically relevant dependence on the cluster size, yet we find a striking and repetitive dependence on the antibody global density. Furthermore, all the clusters produced a degree of degranulation similar to that produced by continuous antibody arrays for each antibody dilution. This confirms the previous observation that for the probed range of cluster size and arrangement, the global density of the antibody predominantly regulates the activation in these systems.

**Figure 2:**
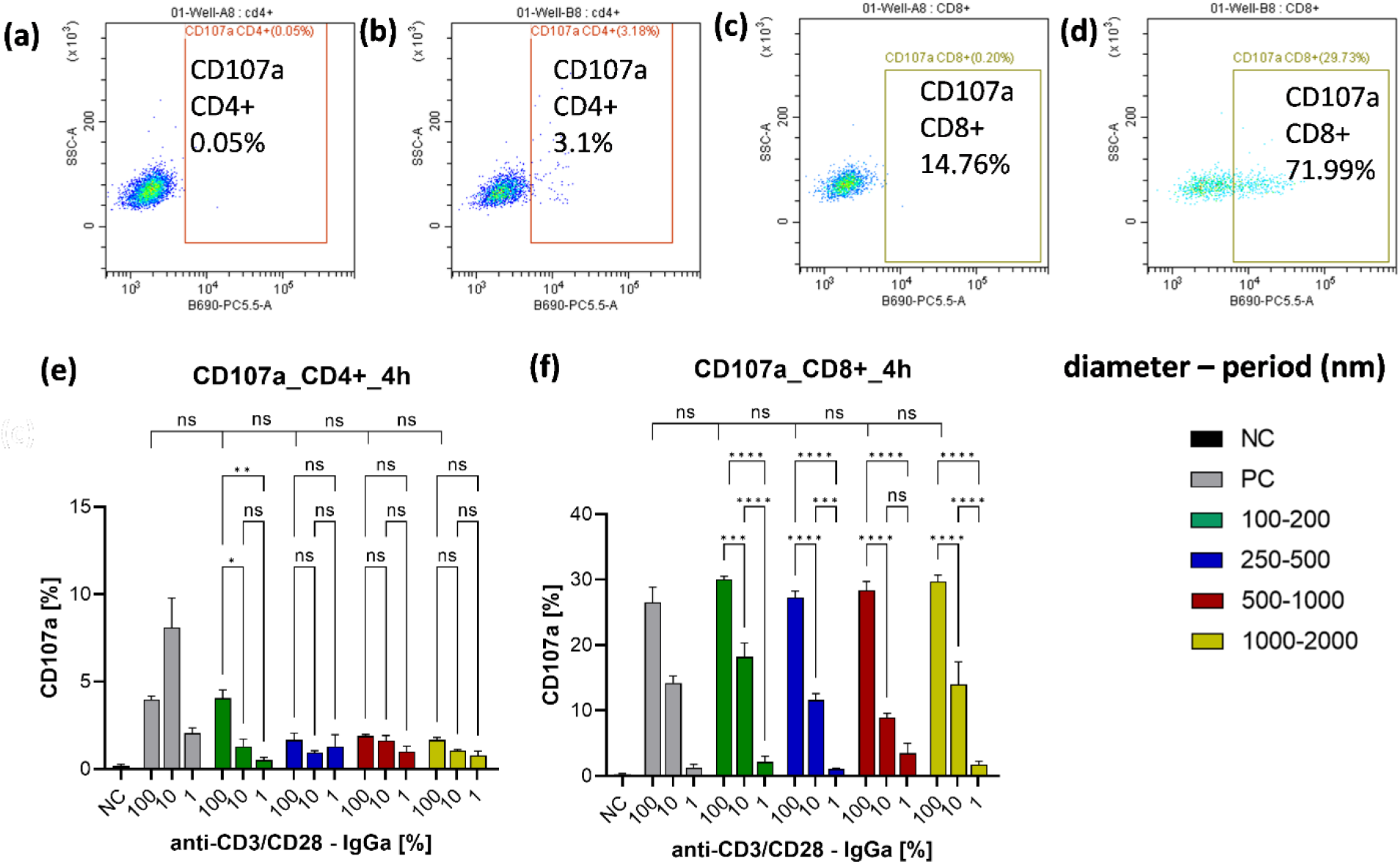
Activation of T cells measured through CD107a expression levels. (a) and (b) FC analysis of CD107a expression by CD4+ T cells stimulated on negative control and 100 nm clusters with no dilution, respectively. (c) and (d) FC analysis of CD107a expression by CD8+ T cells stimulated on negative control and 100 nm clusters with no dilution, respectively. (e) and (f) CD107a expression for CD4+ T cells and CD8+ T cells stimulated on various antibody clusters and dilutions. NC and PC stand for negative and positive controls, respectively. The statistical analysis was performed with Tukey’s multiple-comparison tests using the GraphPad Prism software. * *p <* 0.05, ** *p <* 0.01, *** *p <* 0.001, **** *p <* 0.0001, ns: not significant.

## Membrane fluctuation model for immune cell activation

The obtained correlation between the antibody density and T cell activation, as well as the lack of sensitivity to the probed cluster sizes could be rationalized through the so-called Membrane Fluctuation Model (MFM). The latter was recently conceived by us to predict the response of natural killer cells to the spatial organization and clustering of the activating ligands^25,31^, but its principles are transferable to other cell types. In essence, MFM captures the dynamic coupling between the ligand-receptor binding and the activity of the cell. Since the receptors are embedded within the T cell membrane, their binding and unbinding directly impact the membrane configuration above the cluster (and, due to membrane rigidity, also in the regions between the clusters). On the other hand, the binding and unbinding rates (*K*on, *K*off, respectively) are explicit functions of this average height and the fluctuation amplitude,^39^ and typically, the height and amplitude of fluctuation are smaller in the vicinity of a bond than those of the unbound membrane.^36^ As a result, the probability to bind in the vicinity of an existing bond is strongly enhanced, and the affinity of the ensemble of bonds is increased.^36^ Formation of bonds triggers the signaling and initiates the activation process, which, in turn, changes the state of the cell, enhancing the cytoskeleton dynamics, cytoplasmic flows and/or ion channel activity, among other processes. The ensemble of these processes boosts the dynamics of the membrane, which is reflected in increasing fluctuation amplitudes with activation.^25,39^ In turn, such change in membrane fluctuations acts as a mechanical feedback and leads to changes in the binding and unbinding rates. Thus, depending on the geometry, binding and cooperative effects between nearby receptors on the membrane are affected. If these correlations promote further binding, the amplification process ensues until the number of bonds and the activity of the cell saturates in new steady state.

Mathematically, the model assumes that the activation is proportional to the density 𝒩 of ligand– receptor bonds of the cell, which in the stationary state is given by 𝒩 = 𝜚 *K*_on_/(*K*_on_ + *K*_off_) where 𝜚, *K*_on_, *K*_off_ are the ligand density, and binding and unbinding rates, respectively. The two rates are calculated directly from the first principles describing the ligand–receptor pair as a spring with defined stiffness and rest length (see SI for details). We note that the approximations used within the current version of MFM may underestimate the effect of correlations. Qualitative agreement between the simulation and experimental results is obtained by relating the activity of the cell to the fraction of occupied antibodies within the pattern (𝒫 = 𝒩/𝜚) (see SI for details). This is an estimate that could be ameliorated in future work.

The MFM was applied to experimentally studied geometries in order to calculate the fraction of bound antibodies for the probed cluster configurations and dilutions, separately with and without the feedback loop (Figure 3(a) and (b), respectively). The results clearly show that the activating and costimulatory signal produced by T cell increases upon increasing the global density of antibodies (and thus increasing the number of antibodies available to the cell), regardless of the cluster configuration. Furthermore, the model shows that, for a fixed global density, increasing size of the clusters of antibodies leads to an increase of the activation, although this effect is very weak. This demonstrates that receptor is more likely to bind and rebind its specific ligand near an already-formed ligand–receptor complex due to the membrane-mediated cooperativity.

**Figure 3:**
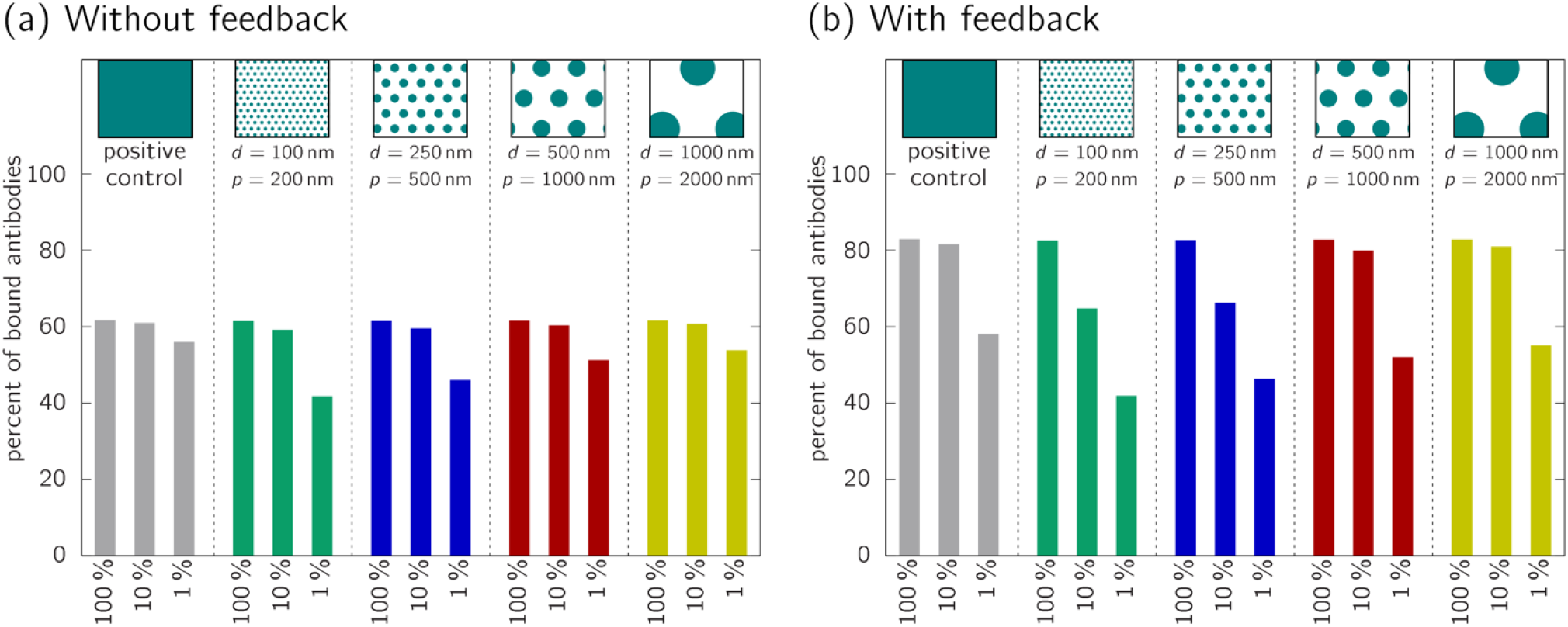
Percentage of bound antibodies upon interacting with the experimentally defined patterns (top row) and antibody densities predicted by the Membrane Fluctuation Model. (a) No mechanical feedback is applied in the calculation. (b) Steady state activity upon amplification.

Moreover, the model data obtained with the feedback loop seems to agree better with the experimental results than the results obtained without the feedback. The lowest probed density of the antibodies, which corresponds to 1% dilution, produced the lowest percentage of bound receptors, which seems insufficient to induce activation. Yet, for the arrays with 10% dilution and arrays with no dilution (100%), the number of bound antibodies grows with the antibody density until it reaches saturation, regardless of the size of the cluster within which the antibodies are confined. This suggests that activation is indeed a process in which the dynamic evolution of the number of formed bonds is influencing the state of the cell and consequently, the state of the membrane.

The model shows cooperative effects of the overall ligand density and clustering for undiluted and 10%-diluted systems, with feedback loop. Yet, this was not observed in the experimental data. This discrepancy between the theory and experimental data has two possible explanations that do not preclude each other. First, the model was parametrized to have the largest sensitivity to patterns at intermediate surface coverage, which is reflected in the 10%-diluted systems (see SI for details). In the experimental data, this effect could vanish due to the large spreading of the experimental data points. Furthermore, we do not measure directly the number of formed bonds, but provide the indirect benchmark of this number, which is the functional response of T cells. This response, in turn, is affected by many additional physiological factors, which likely diminish the relatively weak effect of the ligands clustering.

We, furthermore, conclude that the almost evenly high activation of T cells on all the arrays with undiluted antibodies likely stems from the fact these arrays provide an excess amount of activating and costimulatory stimuli. Namely, even though for antibody-based activation the number of needed activating receptors is supposed to be higher than a few to a few hundred expected for activation induced by natural ligands^32^, the geometry of our patterns suggest that a T cell is exposed to a total of ∼700 000 antibodies. This stems from estimation that a T cell membrane is establishing a contact area with the array of a diameter of ∼10 μm and assuming a dense packing of antibodies with the footprint of about 5 nm × 5 nm within a cluster. Consequently, each cluster of a diameter 100 nm contains ∼300 antibodies in total, and global density is ∼9000 antibodies per μm^2^. Even with relatively low on–off binding rate of the antibody–receptor complex (which typically precludes serial receptor triggering), the MFM suggests that such dense packing of antibodies within the cluster will strongly stabilize the bonds to the receptors. Consequently, a very large fraction of receptors on the cell will be engaged, saturating the signal. On the other hand, assuming that the activating, costimulatory, and mock antibodies are evenly distributed, for 1% antibody dilution, each 100 nm cluster contains, on average, only a few activating or costimulatory antibodies. Consequently, the antibodies are too far apart to benefit from membrane-induced correlations, which precludes triggering of the feedback loop mechanism to enhance the activation.

## Activation by clusters of tightly packed antibodies on surfaces with typically insufficient global density

The results presented so far suggest that grouping activating and costimulatory antibodies into dense clusters, even at global densities that are typically too low to promote T cell activation, could produce the proximity-induced feedback loop. Consequently, tight clustering could promote binding and the triggering of the feedback loop, and ultimately induce the activation of T cells, even at global densities that are below activation threshold. To test this hypothesis, we prepare a new design of an array, denoted as 100 nm/2000 nm/100% configuration (Fig. 4a), that combines (*i*) global density equal to that in the previously studied arrays with 1% dilution and (*ii*) the densest possible packing of activating and costimulatory antibodies within small clusters. Specifically, the new pattern consisted of circular clusters of a 100 nm diameter densely packed with anti-CD3/anti-CD28, hexagonally arranged with 2 μm periodicity (see Fig. 4a). This new 100 nm/2000 nm/100% system was fabricated on Si substrates covered with Ti/Au (1nm/ 4nm), using electron-beam lithography, electron-beam evaporation of silicon dioxide, and liftoff, to allow using the same functionalization chemistry as in the previously described arrays (Fig. S4 shows SEM of the array). Functionalization site selectivity of these arrays was verified using by fluorescence (see Fig. 4c), while their performance was assessed by comparing the state of T-cells to activation on 100 nm/200 nm/1% patterns, where basically no T cell activation was observed following CD69 and CD107a expression (Figs 2 and 3).

**Figure 4:**
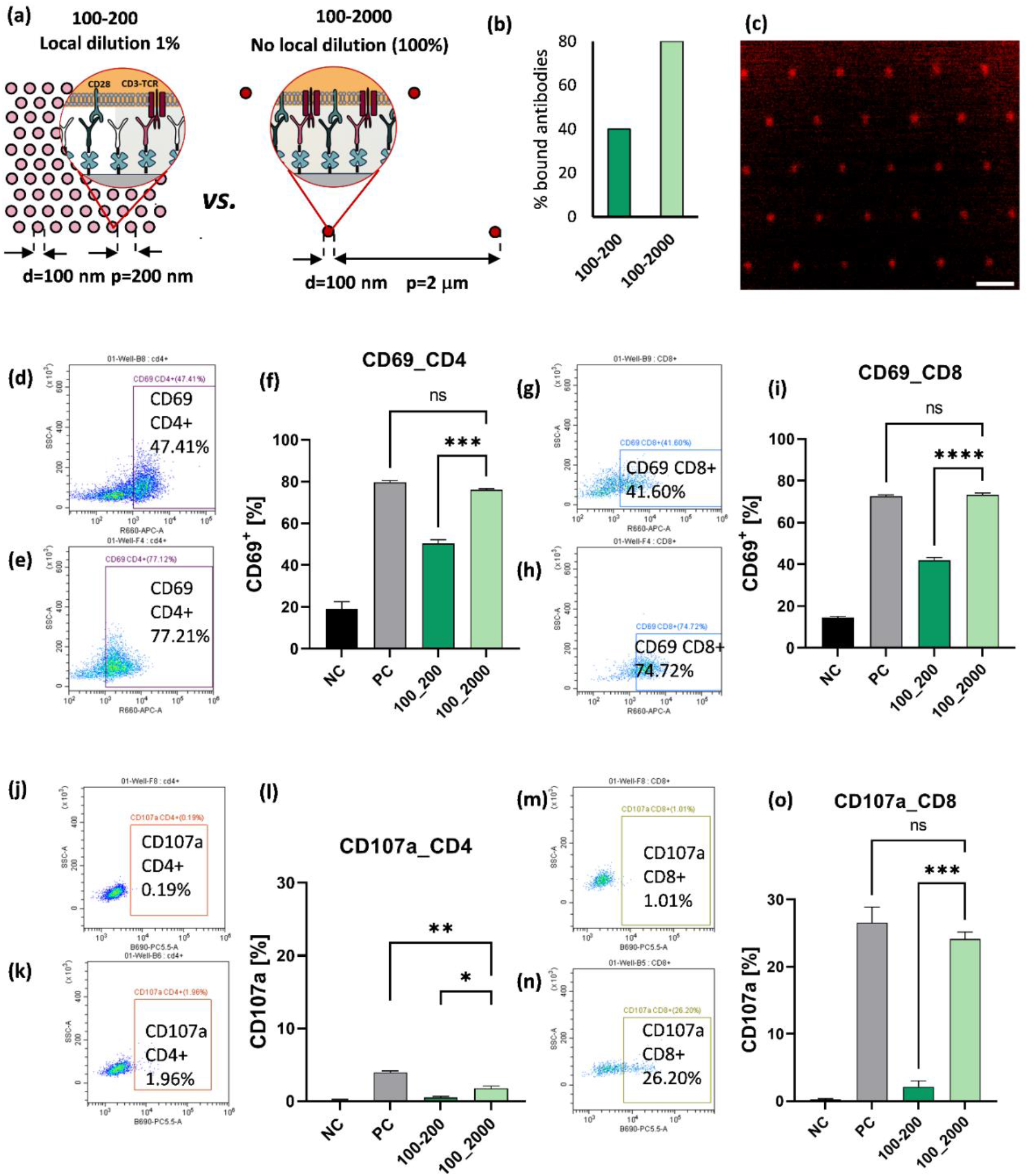
(a) Schematic plot of two nano-arrays with the same theoretical global antibody density and cluster size, but different antibody dilutions within the clusters—1% and 100% (no dilution), and different cluster spacing 200 nm and 2 μm. (b) MFM-based analysis of bound antibodies for two arrays with the same global density: 100 nm/200 nm/1% and 100 nm/2000 nm/100%. (c) Fluorescent image of 100 nm antibody clusters separated by 2 μm. (d) and (e) FC analysis of CD69 expression by CD4+ T cells stimulated on diluted and undiluted clusters with the same global antibody density, respectively. (f) Percent of CD69 positive CD4+ on the tested clusters and control experiments. (g) and (h) FC analysis of CD69 expression by CD8+ T cells stimulated on diluted and undiluted clusters with the same global antibody density, respectively. (i) Percent of CD69 positive CD8+ on the tested clusters and control experiments. (k) and (l) FC analysis of CD107a expression by CD4+ T cells stimulated on diluted and undiluted clusters with the same global antibody density, respectively. (m) Percent of CD107a positive CD4+ on the tested clusters and control experiments. (n) and (o) FC analysis of CD107a expression by CD8+ T cells stimulated on diluted and undiluted clusters with the same global antibody density, respectively. (p) Percent of CD107a positive CD8+ on the tested clusters and control experiments. The statistical analysis was performed with Tukey’s multiple-comparison tests using the GraphPad Prism software. * *p <* 0.05, ** *p <* 0.01, *** *p <* 0.001, **** *p <* 0.0001, ns: not significant.

According to the MFM, as presented in (Fig. 4b), the tight clusters of 100 nm/2000 nm/100% pattern would double the number of formed bonds due to proximity effects and allow for the full T cell activation (light green bar). For the same global density, the 100 nm/200 nm/1% produces an insufficient number of bonds to trigger the mechanical feedback (dark green bar).

Consistently with this prediction, we found that the CD69 expression was substantially higher for the T cells activated on 100 nm/2000 nm/100% than that on 100 nm/200 nm/1%. We also found that the difference between them is about two-fold for both CD4 and CD8 cells, which is very similar to the difference obtained for the calculated number of bonded antibodies. The difference between patterns 100 nm/2000 nm/100% and 100 nm/200 nm/1% becomes even stronger when it comes to the degranulation marker CD107a. For CD8 T cells, this difference is an order of magnitude. Even for CD4+, whose only very small part is cytotoxic, reflecting thus in the overall low numbers of CD107a positive cells, the system 100 nm/2000 nm/100% produced substantially higher marker signal than 100 nm/200 nm/1%. This can be considered as a convincing experimental confirmation that the number of formed receptor–antibody pairs strongly correlates with the degree of activation, but also that correlations between binding events within dense clusters play an important role for the stabilization of these bonds.

Another intriguing fact is that the pattern 100 nm/2000 nm/100% produced as high activation as the positive control surface with continuous dense antibody coverage. The main outcome of these findings is that, while clustering of activating and costimulatory antibodies does not affect T cell activation at all for excessive global antibody density, it does so for low density. Therefore, even though a certain global antibody density could be insufficient for T cell activation if the antibodies are randomly distributed over the surface, the same density can be sufficient if the antibodies are grouped into clusters. In this scenario, the receptor nanoclustering driven by the antibody arrangement compensates for the insufficient antibody density. MFM shows that this phenomenon stems from the fact that the binding of one receptor to an extracellular molecule (ligand or antibody) greatly affects the binding of other receptors in its lateral proximity. These results comply with previous physical modeling, which suggested that, away from a bond, the distance between the T cell and a target cell membrane is set by the size of large phosphatase molecules, which act as a repelling agent keeping the separation between the T-cell membrane and the target surface large^33–35^, while the binding of a relatively small TCR receptor to its ligands produces a close transversal proximity between the T cell and a target cell.

The region of a close transversal proximity laterally extends to ∼40–100 nm around the point of the receptor binding,^36,37^ due to the finite lateral correlation length of the membrane^38^. This further facilitates the formation of more similarly sized ligand–receptor bonds around already formed bonds^39^. The accumulation of these bonds can counteract the repulsive forces between the membranes, further stabilizing the bonds, even in an activated state^40^.

## Summary

Currently, simplistic platforms such as antibody-coated beads use an uncontrolled surface distribution of activating and costimulatory antibodies, delivering the maximum possible dosage of activating stimuli. However, it is increasingly evident that an excess of activation stimuli does not necessarily result in the best antitumor potency of immunotherapeutic T cells.^38^ Moreover, over-activation of T cells often leads to rapid exhaustion, recusing their anti-tumor activity. Therefore, the dosage of these stimuli must be optimized for the clinical context. On the more fundamental level, the control of the activation of immune cells including T cells, clearly involves a response to the mechanical properties of the environment. Still, understanding the interplay between this mechanical and biochemical signals has proved to be a major challenge.

We believe that our experimental findings, rationalized by the membrane fluctuation model, clearly establish an important link between the cell mechanics and activity. This allows us to resolve a long-standing question about the role of patterning in T-cell activation. We demonstrate that patterning plays no role if the T cells is exposed to an abundance of activating ligands. However, for low global concentrations, where sparse organization of ligands is not capable of producing a relevant signal for the cell; dense, small clusters of ligands can trigger the feedback mechanisms and induce the full T cell activation. This shows that, indeed, clustering can be used for the regulation of T cell activation. In the current work, the clusters were pre-defined, allowing to demonstrate that membrane mediated forces play an important role. In the interaction of T cells with target cells, these effects should promote the spontaneous formation of the nanodomains that can trigger the activation process. These results, thus offer significant insights into the fundamental physical mechanisms of T cell activation.

In the more applied setting, our findings have the potential to greatly enhance the rational design of future *ex vivo* T cell activation platforms for immunotherapy. By further exploring the strategy of tuning activation by antibody clustering, one could design platforms that balance adequate activation to ensure a high level of CAR transfection and proliferation, achieving clinically sufficient T cell quantities, while minimizing exhaustion. This balance may promote differentiation into the immunotherapeutically relevant central memory T cell phenotype, ultimately ensuring high antitumor potency.

Further work, both from the experimental and modeling side is, however, required to fully understand the interplay between T cell mechanics and signaling. This calls for further usage of reductionist platforms,^41^ where the mechanical properties of the environment can be strictly controlled and lymphocyte cell activation can be studied on different length scales^22,42–45^. This includes recent nanotechnological advances that now allow for the control of the position of ligands at the individual molecule level,^46–48^ which can be combined with varying substrate elasticity,^49–52^ and nanoscale topography of non-deformable,^53^ or deformable^54,55^ setting to measure forces involved in the activation process ^56,57^. Studying T cell activation of such platforms that combine several stimuli should allow for deepening our understanding of their combined effects on T cell activation, a task that we aim to address in future work.

## Materials and Methods

### Fabrication of the arrays

Dense monolayers of 200 nm polystyrene nanoparticles were fabricated to Silicon substrates using a previously reported method. Briefly, polystyrene nanoparticles (Polyscience Inc) were suspended in 2:1 mixture of water:ethanol, and the solution was slowly dispensed from automatic syringe on the surface of Langmuir–Blodgett through filled with DI water, using a glass skinned positioned on the through barrier at an angle^58^. Larger polystyrene nanoparticles were assembled by mechanical rubbing of particle powder between two PDMS surfaces, and the particle monolayer formed on PDMS was mechanically transferred to Silicon using PEI adhesive film^27^. The assembled particles were trimmed to half of their diameter using Oxygen plasma etching. Then, a thin film of Ti/Au (1 nm/5 nm) was deposited through the formed polystyrene mask by electron beam evaporation, and liftoff in hot chlorobenzene removed the particle, leaving silicon disks surrounded by Au. For electron-beam patterned arrays, silicon substrates were first coated with Ti/Au (1 nm/5 nm) by electron beam evaporation, and then by a film of positive e-beam resist (PMMA, 950 K). The films were then exposed by electron-beam writing tool (EBPG 5150), and developed for 1 minute in MIBK:IPA 1:3 solution. The resulting surfaces consist of arrays of silicon nanodisks surrounded by a gold background.

### Biofunctionalization of the arrays

Prior to functionalization, samples were cut into 1×1 cm and were fixed to the underside of a bottomless 96-well plate using PDMS as sealant. In this way, cells could be directly incubated over the samples and would interact solely with the functionalized arrays. The plates were then treated by UV-ozone (Novascan) to activate the native oxide layer of the silicon nanodisks, which were subsequently functionalized using 3-aminopropyl triethoxysilane (APTES, Merck). For this, samples were incubated 30 min in a 5% ethanolic solution of APTES, followed by 30 min baking at 60°C. The amine terminated nanodisks were then derivatized with biotin by soaking overnight in a 1 mM aqueous solution of biotin N-hydroxysuccinimide ester (NHS-Biotin, ThermoFisher). To minimize non-specific interactions, the gold background was passivated using a 0.5 mM ethanolic solution of mercapto polyethylene glycol (thiol-PEG3400, Nanocs). The plates underwent a final sterilization step by 1 hour soaking in ethanol and all ensuing steps were performed in a sterile laminar flow using sterile buffers. Green-fluorescent avidin (Neutravidin Oregon Green 488, ThermoFisher), was coupled to the biotin terminated nanodisks by incubation for 90 min at a concentration of 25 μg/mL in phosphate buffered saline (PBS) with 2% bovine serum albumin (BSA). The plates were thoroughly rinsed with PBS containing 0.1% Tween20 (PBST), and then with PBS only. Finally, the samples were biofunctionalized with antibodies by 90 min incubation in a 1:1 mix of biotinylated anti-human CD3 (OKT3 clone) and anti-human CD28 (both from Biolegend) at a concentration of 2 μg/mL in PSB with 2% BSA. Finally, the samples were consecutively washed with PBST and PBS and were directly used for cell experiments.

### PBMC isolation and activation

Peripheral blood mononuclear cells (PBMCs) were isolated from blood using the FICOL gradient. First, blood was diluted with PBS augmented with 2% fetal bovine serum (FBS), at a 1:1 ratio, then loaded on FICOL gradient, and centrifuged at 16°C at 1200 g (with no breaks or acceleration). The PBMCs were collected as the middle disc and a small portion of the underlying phase but taking care not to withdraw the pellet, washed three times with at least 1:2 with PBS 2% FBS at room temperature, and sedimented at 500 g. The cells were finally suspended in the final medium in the ratio of 2 mL per 7 mL of collected blood, counted, and diluted with the medium to final concentration of 1 × 10^6^ cells per mL. The cells were then seeded onto the surfaces in the growth medium containing <2% serum and 50 units of IL-2 and left inside incubator to adhere for up to 18 h.

### Flow cytometry

For flow cytometry measurements, 50 000 cells were used per well. The cells were stained with respective fluorophore-conjugated antibodies at 1:1000 dilution and incubated for 30 minutes on ice. Thereafter, the cells were washed, and the dead cells were stained with DAPI (1:1000 in PBS). All the samples were analyzed on a CytoFLEX LX flow cytometer (Beckman Coulter). For analysis, the fraction of CD3-positive cells was calculated, and CD3-positive cells were then analyzed for staining with the other antibodies employed for staining. The antibodies used for staining were PE anti-human CD3, FITC anti-human CD4, APC Fire 750 anti-human CD8, APC anti-human CD69, and PerCP-Cy5.5 anti-human CD107a (all from Biolegend).

### Fluorescence and scanning electron Imaging

Two sets of samples with nanoarrays were biofunctionalized as detailed previously without being mounted in 96-well plates. Instead, they were placed in 24-well plates into which PBMCs were seeded. After incubation overnight, cells were fixed in 4% paraformaldehyde at 4 °C for 15 min. One set was used for fluorescent microscopy while the other was dedicated to scanning electron microscopy. For fluorescence, the cells were stained for cytoskeleton using AlexaFluor 555 Phalloidin (ThermoFisher), mounted with DAPI containing medium (Abcam), and imaged with a Zeiss LSM 880 confocal microscope. For the other set, the fixed cells were treated with three successive dips in absolute ethanol for 5 min each, then dried in a critical point dryer, sputtered with a thin layer of gold and imaged using a Verios 460L HRSEM.

### Statistics

For the flow cytometry experiments, each condition was prepared in triplicate and cells in each individual well were measured. For gating, antibody-positive population were set against the negative control which consisted of naïve PBMCs cultured in standard conditions. Results over the triplicates were then averaged. Statistical analyses were performed using Prism (GraphPad), where results were considered significant for p<0.05.

## Acknowledgements

A.-S.S. and M.S. gratefully acknowledge the support of the German Research Foundation via Joint Project SM 289/10-1. M.S. acknowledges the support of the Israel Science Foundation Project 2016/21. A.-S.S. and P.N. received further support by the German Science Foundation and the French National Research Agency Project SM 289/8-1.

