## Supplementary Information for "Deciphering the role of the nanoscale clustering of activating and costimulatory antibodies on T cell activation"

### Supporting experimental details

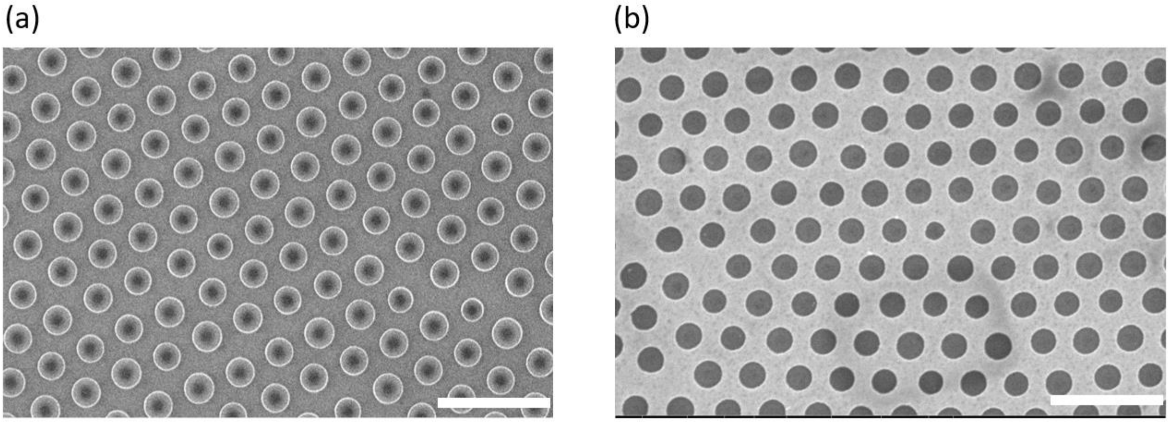


**Figure S1**: Example of the fabrication results: (a) Array of polystyrene nanospheres with the original diameter of 500 nm, after trimming by Oxygen plasma to half of their original diameter. (b) Array of Silicon disks surrounded by Au mesh obtained after Au deposition and liftoff. Scale bar: 1 μm

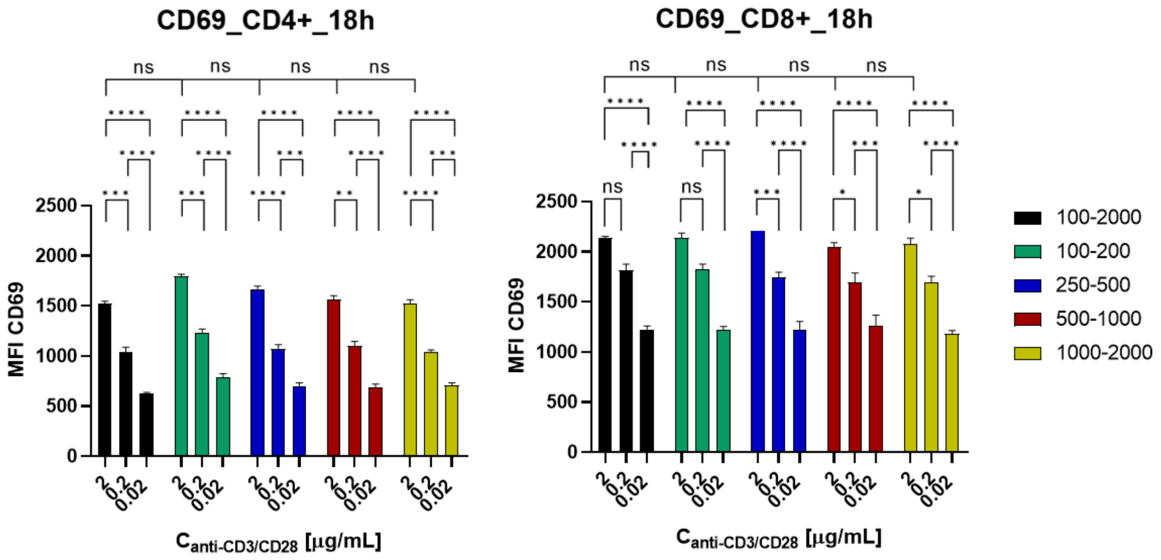


**Figure S2**: Mean Fluorescence Intensity (MFI) of CD69 expression for CD4+ T cells and CD8+ T cells stimulated on various antibody clusters and dilutions, as obtained by flow cytometry. NC and PC stand for negative and positive controls, respectively. The statistical analysis was performed with Tukey's multiple-comparison tests using the GraphPad Prism software. $* p < 0.05$, $** p < 0.01$, $*** p < 0.001$, $**** p < 0.0001$, ns: not significant.

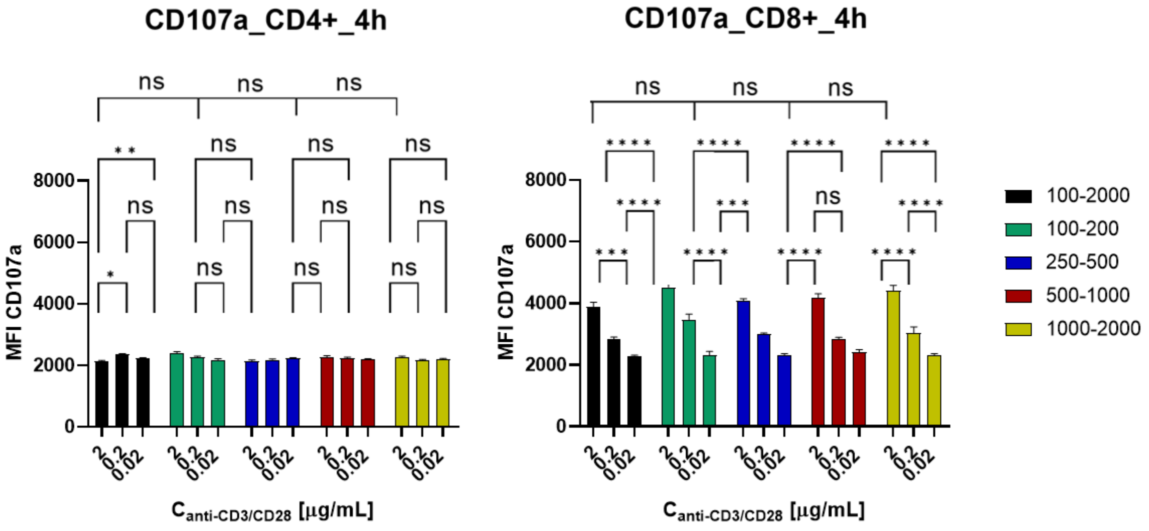


**Figure S3**: Mean Fluorescence Intensity (MFI) of CD107a expression for CD4+ T cells and CD8+ T cells stimulated on various antibody clusters and dilutions, as obtained by flow cytometry. NC and PC stand for negative and positive controls, respectively. The statistical analysis was performed with Tukey's multiple-comparison tests using the GraphPad Prism software. $* p < 0.05$, $** p < 0.01$, $*** p < 0.001$, $**** p < 0.0001$, ns: not significant.

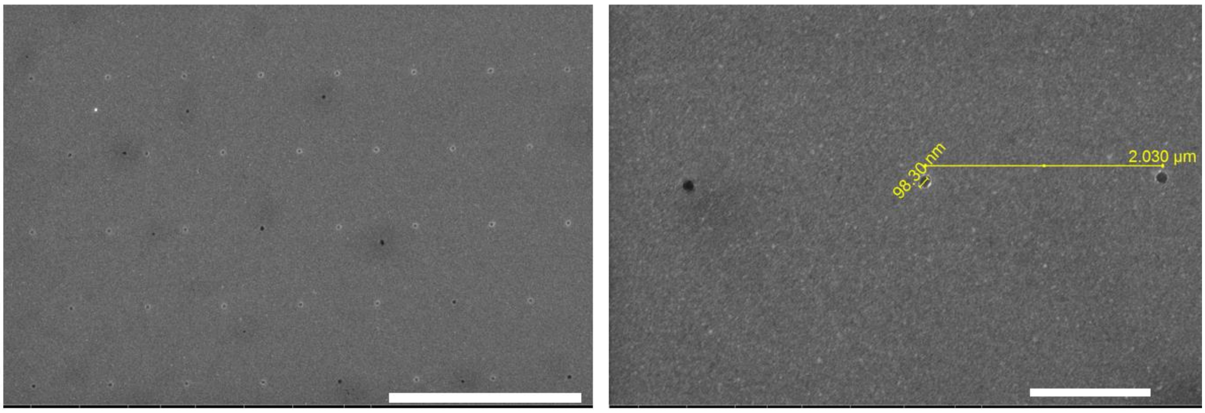


**Figure S4**: SEM of 100 nm Silicon Dioxide clusters arranged with the spacing of 2 μm. Left: X 8K magnification, scale bar 5 μm, Right: X 25K , scale bar 1 μm.

**Membrane fluctuation model**

In order to explain the results of experimental measurements we have made calculations using membrane fluctuation model (MFM). This model has already been successfully applied to explain activation of NK cells [see ref. 25 in the main text]. In MFM, it is assumed that there is a feedback loop between the number of bonded receptors and the activity of the cell: increasing the number of bindings is increasing activity of the cell which manifest itself by a growth of fluctuations of the membrane, which, in turn, promotes the formation of further bonds.

We describe the membrane by its height over the flat substrate $h(\boldsymbol{r})$, where $\boldsymbol{r}$ is a two-dimensional vector defining a point on the substrate plane. We assume that the energy of each configuration of the membrane is given by a Hamiltonian

$$\mathcal{H}_{\text{membrane}} = \int_{\mathcal{D}} \mathrm{d}\boldsymbol{r} \left[\frac{\kappa}{2}\left(\nabla^2 h(\boldsymbol{r})\right)^2 + \frac{\gamma}{2}(h(\boldsymbol{r}) - h_0)^2\right] + \frac{\lambda}{2}\sum_{j=1}^{M}\left[h(\boldsymbol{r}_j) - \ell\right]^2, \quad \text{(S1)}$$

where $\kappa$ is the bending stiffness of the membrane, $\gamma$ is the strength of unspecific potential keeping the unbound membrane at the average transversal distance from the patterned surface $h_0$. The second term in the Hamiltonian (S1) describes interaction of the membrane with $M$ antibodies bound to receptors in positions $\boldsymbol{r}_j$ for $j = 1,2,3, \dots, M$. Finally, $\ell$ is the rest length of the antibody-receptor bond that has a stiffness $\lambda$. The exact location of antibodies, as well as exact size of a rectangular domain $\mathcal{D}$ of integration, is defined by the pattern on the substrate, as discussed below.

In the MFM, the amplitude of fluctuation of the membrane of an inactive cell $\sigma^{\text{inactive}}$ is increased by the amplitude $\sigma^{\text{active}}$ of fluctuations associated with the activity of the cell. For an inactive cell, which is not bound to antibodies, the membrane hovers at an average transversal distance $h_0$ from the patterned surface with the Gaussian fluctuations with a standard deviation $\sigma^{\text{inactive}} = \langle (h(\boldsymbol{r}) - h_0)^2 \rangle^{1/2} = (64\kappa\gamma)^{-1/4}$.

Upon binding, the activity of the cell may change, giving rise to another component of fluctuations $\sigma^{\text{active}}$ which could enhance the dynamics of the membrane. MFM assumes the relation between $\sigma^{\text{active}}$ and the activity $\mathcal{P}$ in a form of a logistic curve:

$$\sigma^{\text{active}}(\mathcal{P}) = \frac{1}{2}\sigma_{\max}\left[\tanh\left(\frac{\mathcal{P} - \mathcal{P}_0}{\mathcal{P}_{\mathrm{w}}}\right) + 1\right]. \quad \text{(S2)}$$

Here, the parameters of the model are $\sigma_{\max}$ which denotes the maximal amplitude of active fluctuations, $\mathcal{P}_0$ is the middle value of activity, and $\mathcal{P}_{\mathrm{w}}$ is the width of the sigmoid curve.

Typically, upon the formation of first bods, a laterally dependent height and fluctuation profile develops (see ref. 36 of the main manuscript for details). This means that, in reality, every antibody experiences a different average height and fluctuation amplitude $\sigma_j$, depending on the evolution of the bonds in its vicinity, yielding correlations between binding events. In MFM the fluctuations of the membrane are assumed to be a convolution of inactive and active fluctuations, and hence, total fluctuation amplitude of the membrane at the position of the antibody is

$$\sigma_j = \left[\left(\sigma_j^{\mathrm{inactive}}\right)^2 + (\sigma^{\mathrm{active}})^2\right]^{1/2}. \tag{S3}$$

For a given pattern of bound antibodies average height and fluctuation amplitude $\sigma_j^{\mathrm{inactive}}$ can be calculated from Hamiltonian (S1) using the method of path integral.

The fluctuation amplitude and the transversal distance of the membrane from the patterned array both explicitly enter the rates (see ref. 38 of the manuscript). Consequently, one should calculate binding and unbinding rates $K_{\mathrm{on}}$ and $K_{\mathrm{off}}$ rates for each antibody $j$. Notably, these rates should carry the information of the likelihood of finding a receptor in the binding range of the antibody.

Finally, by knowing the rates, one can calculate the activity of the cell, which is assumed to be related to the number of formed bonds $\mathcal{N}$ in stationary state given by

$$\mathcal{N} = \varrho\overline{K_{\mathrm{on}}}/(\overline{K_{\mathrm{off}}} + \overline{K_{\mathrm{on}}}). \tag{S4}$$

Here, $\varrho$ is the total density of antibodies, and $\overline{K_{\mathrm{on}}}$ and $\overline{K_{\mathrm{off}}}$ denote average binding and unbinding rates, respectively. We approximate $\overline{K_{\mathrm{on}}}$ and $\overline{K_{\mathrm{off}}}$ as a sum of all $K_{\mathrm{on}}$ and $K_{\mathrm{off}}$ rates calculated for all antibodies within a unit area. However, because of the large number of configurations, calculating these averages is very challenging. We therefore, estimate the average rates by considering only the most probable scenarios: $\overline{K_{\mathrm{on}}}$ is calculated as a rate of binding of the receptors to an antibody from a fully unbound membrane, and $\overline{K_{\mathrm{off}}}$ is calculated as the rate of bond breaking in a fully bound cluster.

Naturally, this approach provides only an approximation of the correlations in the system. Furthermore, we sent the likelihood of finding a receptor above the antibody to unity. To compensate for these approximations, we define the activity of the cell as $\mathcal{P} = \mathcal{N}/\varrho$:

$$\mathcal{P} = \overline{K_{\mathrm{on}}}/(\overline{K_{\mathrm{off}}} + \overline{K_{\mathrm{on}}}). \tag{S5}$$

Consequently, $\sigma^{\mathrm{active}}$ is enhanced for low densities of antibodies where the collective binding may be of importance, and reduced for larger densities where the limited number of receptors is relevant. This is clearly different from the original application of MFM [ref. 25 in the main text] (where the activity of cell was related to $\mathcal{N}$), in which the densities of ligands on the substrate were not exceeding 250 ligands/μm$^2$ and were much lower than densities of up to 9000 antibodies/μm$^2$ present in the current experiment for undiluted antibodies. This guaranteed larger distances between ligands, smaller total densities of bonds, and the number of receptors available on the cell membrane was not a limiting factor. As a result, both of the effects described above were not relevant, and explicit density dependence was maintained. Notably, for our calculations of cell activity at low densities (1% dilution, low and high level of clustering), both approaches provide the same conclusions, as the overall density $\varrho$ is identical in systems that are being compared.

In the calculations without a feedback, we assume the fluctuations to be only inactive by taking $\sigma^{\mathrm{active}} = 0$, regardless of the value of $\mathcal{P}$. For the calculations with the feedback loop we calculate $\sigma^{\mathrm{active}}$ from the sigmoid relation (S2) and increase the fluctuations of the membrane following Eq. (S3). This yields new $\overline{K_{\mathrm{off}}}$ and $\overline{K_{\mathrm{on}}}$, and update in the cell activity $\mathcal{P}$, and consequently a new $\sigma^{\mathrm{active}}$, until the steady state is achieved. The results are presented in Fig. 3b and Fig. 4b of the main text.

The sigmoid curve (S2) has been fitted graphically to make the feedback loop mechanism work most efficient for moderate densities of antibodies (dilution 10%). While this naturally, overemphasizes the correlations in this range of parameters, as seen in experiments, it provides a good response at the density extremes (no dilution and dilution to 1%), as argued above. The numerical values of all the parameters used in our calculations are presented in Table S1.

**Table S1**: Numerical values of parameters used in calculations.

| name of parameter | symbol | value used in calculations |
|---|---|---|
| strength of potential acting on the membrane | $\gamma$ | $4 \times 10^5\ k_\mathrm{B}T/\mu\mathrm{m}^4$ |
| equilibrium height of the membrane | $h_0$ | 80 nm |
| bending stiffness of the membrane | $\kappa$ | $30\ k_\mathrm{B}T$ |
| length of bond coupling | $\ell$ | 40 nm |
| bond spring constant | $\lambda$ | $2.5 \times 10^4\ k_\mathrm{B}T/\mu\mathrm{m}^2$ |
| binding pocked on the receptor | $\alpha$ | 10 nm |
| intrinsic affinity of bond | $\varepsilon_\mathrm{b}$ | $9\ k_\mathrm{B}T$ |
| maximum of active fluctuations | $\sigma_\mathrm{max}$ | 5 nm |
| half fluctuation density | $\mathcal{P}_0$ | 0.715 |
| width of activity curve | $\mathcal{P}_\mathrm{w}$ | 0.19 |
| diameter of antibody | $d_\mathrm{antibody}$ | 5 nm |

### Patterned substrate

In order to model the experimental system of a pattern consisting of circles of the diameter $d$, forming a hexagonal lattice with spacing $p$ with antibodies attached only inside the circles with a dilution $c$, we have used the following protocol: First, we set the system shape $\mathcal{D}$ to be a rectangle with periodic boundary conditions (see top row of Fig. 3 in the main text). We choose the width and height of it to be the smallest possible integer multiple of $p$ and $\frac{\sqrt{3}}{2}p$, respectively, for which both width and height are not smaller than $5\xi_\parallel$, where $\xi_\parallel = (\kappa/\gamma)^{1/4}$ is the membrane correlation length in the direction parallel to the substrate. This way we ensure that the periodic boundary conditions do not affect the calculations significantly. Second, we place circles of diameter $d$ in the domain $\mathcal{D}$ to form a hexagonal lattice. Antibodies are placed on a regular square pattern with a spacing $d_\mathrm{antibody}$ equal to their typical size. In the third step, we remove the antibodies that do not lay inside the circles forming a hexagonal lattice. Finally, we include the dilution $c$ of active antibodies by removing each antibody with a probability $1 - c$. This way we determined the number of antibodies $M$ and their locations $\boldsymbol{r}_j$ within the domain $\mathcal{D}$. These parameters were then used in Hamiltonian (S1) for further calculations as described in the previous subsection.

We note that while for a system with 100% dilution of active antibodies, pattern of antibodies generated by the above method is fully deterministic, for smaller dilutions the exact locations of antibodies are subject to randomness. For this reason, for each system with a dilution $c < 100\%$ we have repeated the calculation of the percentage of bound antibodies several (not less than 10) times. In the main text we present averages of these calculations and, as we have checked, the dispersion of the results was in each case below the resolution of our graphs.

In special case of the positive control (PC) patterns, where the whole substrate is covered by antibodies, we have taken $\mathcal{D}$ to be a square of the size $5\xi_\parallel$ and we have skipped the third step of the protocol of placing antibodies described above.

In order to keep the computational time reasonable, we have decided to treat separately the special pattern with $d = 1000\,\mathrm{nm}$, $p = 2000\,\mathrm{nm}$, and $c = 100\%$. In this case, the distance between circles forming hexagonal pattern exceeds $5\xi_\parallel$. We have, therefore considered an infinite system ($\mathcal{D} = \mathbb{R}^2$) with a single circle of active antibodies of the radius $d$ in the origin, rather than periodic boundary conditions. This greatly reduced the computation time, for the price of ignoring possible correlations on the membrane at distances larger than 10 correlation lengths. According to previous calculations (ref. 36 in the manuscript), this should not affect the obtained results.
